# Cellular and molecular changes in the human osteoarthritic and aging hip pulvinar

**DOI:** 10.1101/2024.04.21.590119

**Authors:** Bahaeddine Tilouche, Stephanie Farhat, Spencer Short, Mariya Somyk, Paul Beaulé, Sasha Carsen, George Grammatopoulos, Daniel L. Coutu

**Affiliations:** Regenerative Medicine Program, Ottawa Hospital Research Institute; Department of Cellular & Molecular Medicine, University of Ottawa; Children’s Hospital of Eastern Ontario; Division of Orthopaedic Surgery, The Ottawa Hospital

**Keywords:** osteoarthritis, sub-synovial adipose tissue, pulvinar, hip, stem cells, mesenchymal progenitors, confocal microscopy, vasculature, innervation

## Abstract

Osteoarthritis (OA) represents a multifaceted pathology characterized by intricate signaling across various joint tissues, where the sub-synovial adipose tissue (ssAT) has been suggested to play diverse roles, from serving as a stem cell reservoir, mechanosensing, serving as a neuroendocrine organ, to modulating inflammation. In this study, we aimed to uncouple the cellular and molecular alterations within the human hip ssAT (the pulvinar) linked to OA and aging, elucidating the distinct contributions of disease onset and progression versus normal aging. Our findings show a pronounced increase in mesenchymal stem/progenitor cells (MSPCs) in the osteoarthritic pulvinar, associated with the upregulation of putative MSPC markers (DPP4, and THY1), indicating an adaptive repair response. Concurrently, in OA patients we observed an altered immune landscape featuring reduced innate immune cells and elevated exhausted CD8+ cells, along with upregulation of genes critical for inflammation and fibroblast activation. Our findings reveal a nuanced picture of OA, where increased stem cell numbers and vascularization, combined with specific gene expression patterns differentiate OA from normal aging. This study not only delineates the roles of inflammation, immune regulation, and stem cell activity in the OA pulvinar but also identifies potential therapeutic targets to modulate these pathways, offering novel insights into OA as a complex interplay of degenerative and intrinsic tissue repair.

---

Osteoarthritis (OA) is a degenerative disease of synovial joints affecting over 13.6% of the population in Canada^1^. Although some cases of OA are idiopathic, most cases are caused by natural aging, obesity, overuse (wear and tear), or congenital malformations of the joints creating impingement^2^. As OA progresses, it results in chronic pain, decreased mobility, and low quality of life^2^. Because of its prevalence and impact on the life of patients, OA is associated with an important socio-economic burden. Current OA treatment involve pain management, physical therapy, and eventually surgery (including joint replacement surgery)^3^. More experimental approaches involve cell therapy (using mesenchymal stromal cells or platelet-rich plasma, for instance)^4^, microfractures, and microabrasion^5^. However, these have yet to demonstrate clinical benefit in large cohorts in well controlled, randomized clinical trials^6^. Therefore, there is an incentive to identify novel therapeutic targets or to develop new approaches for cell-based regenerative therapies for OA treatment.

While the biomechanical causes of OA are well understood, the biological cascade leading to compromised joint function remains unclear. However, recent studies indicate that the pathogenesis of OA involves all synovial joint tissues (synovial membrane^7^, articular cartilage^8^, ligaments^9^, menisci^9^, and subsynovial adipose tissues^2^ [ssAT]), as opposed to articular cartilage alone. Indeed, some studies suggest that infrapatellar ssAT (Hoffa’s fat pad)^10^ in the knee joint is involved in OA pathogenesis and joint tissues homeostasis in general^11,12^, by playing a role as a: 1) stem cell reservoir^13^, 2) mediator of inflammation^14^, 3) mechanosensor/proprioceptor^15^, and 4) neuroendocrine organ^16^. While ssATs are found in most synovial joints, most studies thus far focused on the knee infrapatellar ssAT. In this study we focused on the human hip pulvinar, a large ssAT found in the acetabular fossa. We specifically asked how the stem cell content, immune pathways, and cellular architecture of the pulvinar are altered in OA patients and during aging. We found that the number of mesenchymal stem/progenitor cells is greatly increased in OA patients and that several genes involved in immune-related pathways are dysregulated in OA. Moreover, we found that the osteoarthritic pulvinar showed increased vascularity but that its innervation was not affected. Taken together, our results provide the first detailed analysis of the cellular and molecular changes occurring in the aging and osteoarthritic human pulvinar.

## Results

### Patient’s demographics and experimental design

To uncouple the effect of normal aging and osteoarthritis on the cellular and molecular composition of the hip pulvinar, we recruited patients scheduled for hip arthroscopy or arthroplasty at the Division of Orthopedic Surgery of The Ottawa Hospital or at the Children Hospital of Eastern Ontario. Patients were assigned to one of four experimental groups (G1-G4) based on age and radiographic evidence of osteoarthritis (Fig.1A). After excluding samples that were mishandled during processing or excluded by the surgeons, we obtained a total of 135 samples consisting of arthroscopic biospies of the pulvinar (G1, n=21 and G2, n=59) or the entire pulvinar (G3, n=28 and G4, n=27). The samples were then processed for one of three assays: 1) in vitro analysis of stem cell content, 2) mRNA isolation and analysis, and 3) fixation for immunostaining and confocal imaging (Fig.1B). The larger samples were bisected for use in multiple assays. Patients’ average age was 18.3+/-0.6 for G1, 32.3+/-2.6 for G2, 35.5+/-2.1 for G3, and 78.2+/-2.9 for G4 (Fig.1C). Their average body mass index (BMI) was 26.0+/-2.6 (G1), 25.1+/-1.5 (G2), 28.6+/-2.4 (G3), and 26.9+/-1.8 (G4, Fig.2D). While we aimed for equal numbers of male and female patients, due to recruitment we obtained slightly more females in groups 1 and 2, and slightly more males in groups 3 and 4 (Fig.1E). The arthroscopic biopsy samples were relatively small (0.035g+/-0.016 for G1, 0.029g+/-0.009 for G2) compared to the arthroplasty samples (0.853g+/-0.697 for G3, and 1.551g+/-0.724 for G4) (Fig.1F). Out of the 135 samples obtained, 125 were analyzed so far: 28 were analyzed for G1, 43 for G2, 27 for G3 and 27 for G4 (Fig.1G). Out of these, and accounting for the larger bisected samples, we performed in vitro stem cell assays on 61 samples (13 for G1, 6 for G2, 23 for G3, and 19 for G4), we used 49 samples for RNA isolation (12 each for G1 and G2, 15 for G3, and 10 for G4), and we fixed 57 samples for immunostaining and confocal imaging (17 for G1, 5 for G2, 18 for G3, and 17 for G4) (Fig.1H).

**Figure 1.**
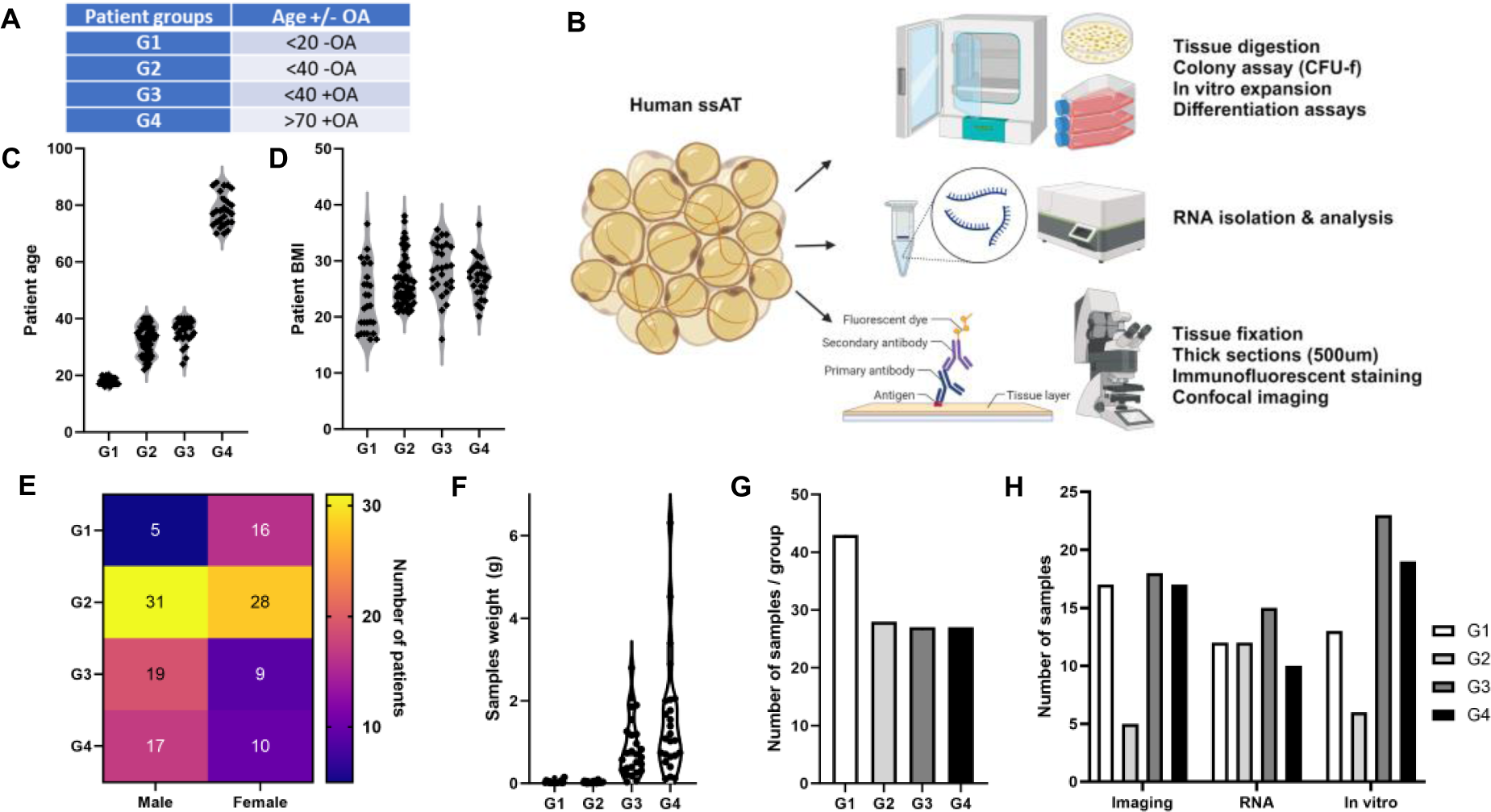
Patients’ demographics and experimental design of the study. A) Patients scheduled for hip arthroscopy or arthroplasty were recruited and assigned to one of four groups (G1-G4) depending on age and radiographic evidence of osteoarthritis. B) Experimental design. ssAT samples obtained from arthroscopic biopsies or arthroplasty were either enzymatically dissociated for in vitro analysis of stem cell content (top), snap frozen for RNA isolation and analysis (middle) or fixed for immunostaining and confocal imaging (bottom). C-D) Distribution of patients per group based on age (C), BMI (D) and sex (E). F-H) Distribution of sample per groups based on weight (fresh samples (F), number of samples obtained per group (G) and sample allocation to various assays (H). For large arthroplasty samples, tissues were bisected for use in multiple assays.

**Figure 2.**
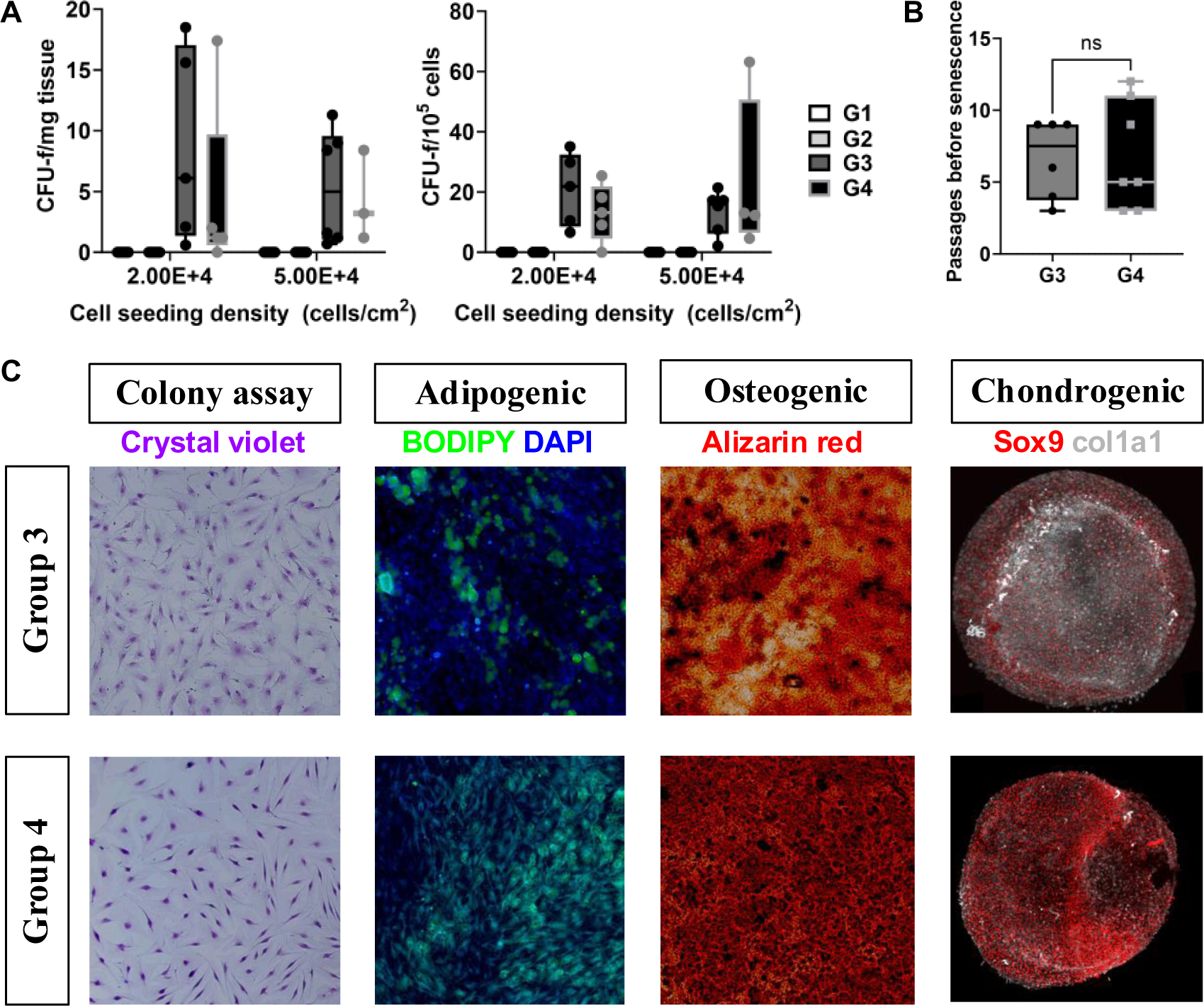
The osteoarthritic pulvinar is a mesenchymal stem/progenitor cells reservoir. Freshly obtained pulvinar tissues were enzymatically dissociated and viable cells were seeded in 24-well plates at clonal density in human Mesencult medium. A) Colony forming unit-fibroblast (CFU-f) assays were performed at various initial seeding densities, but CFU-f’s were only observed starting at 2 and 5×10^4^ cells/cm^2^. An initial seeding density of 5×10^5^ cells/cm^2^ was used in subsequent assays. CFU-f’s were never observed when samples from non-osteoarthritic patients were analyzed (G1 and G2, n=6 and 13 respectively, for all seeding densities tested). B) CFU-f’s from G3 and G4 samples were expanded by passaging at low density until they reached senescence. The primary MSPC cell lines could be expanded for an average of nine passages. C) Multilineage differentiation assays of passage 3 MSPCs obtained from G3 and G4 pulvinars show that each primary cell line tested possessed adipogenic, osteogenic and chondrogenic cells potential (n= 5 and n= 6 respectively for G3 and G4).

### Increased mesenchymal stem/progenitor cell content in the osteoarthritic pulvinar

Mesenchymal stem/progenitor cells (MSPCs) are found in various connective tissues including bone, bone marrow, periosteum, and more. Although their precise anatomical niche, physiological role, and developmental origin remains unclear, they have been proposed to play key roles in tissue regeneration^17^, either by their paracrine effects or differentiation potential^18^. They are normally characterized by their ability to form colonies of adherent cells in vitro, their multilineage differentiation potential into adipocytes, osteoblasts/osteocytes, and chondrocytes, as well as their expression of a few cell surface markers that are non-specific to any cell types in vivo^18^. Since ssATs have been proposed to be a stem cell reservoir^13^ participating in synovial joint tissue homeostatic maintenance, we assessed the MSPC content of the hip pulvinar in our four patient groups using in vitro colony forming assays and multilineage differentiation assays.

Freshly obtained ssAT samples were enzymatically dissociated and viable cells were counted manually or using an automated cell counter. Cells were then cultured in human Mesencult medium (StemCell Technologies) in 24-well plates, in triplicates or more depending on the total number of viable cells obtained. We tested several cell seeding densities but only obtained reliable and clonal colony forming cultures (colony forming unit-fibroblasts, CFU-f’s) at an initial seeding density of 5×10^4^ cells/cm^2^ (Fig.2A). Unexpectedly, we were never able to observe CFU-f’s in cultures derived from non-osteoarthritic patients (G1, n=6; and G2, n=13) at any cell density tested (Fig.2A). With an initial cell seeding density of 5×10^4^ cells/cm^2^, we obtained an average of 5.37+/- 2.36 CFU-f’s per milligram of tissue in G3, equivalent to 13.49+/-3.60 CFU-f’s per 10^5^ viable cells seeded (Fig.2A). Similarly, we observed an average of 4.27+/-1.86 CFU-f’s per milligram of tissue in G4, or 23.28+/-0.25 CFU-f’s per 10^5^ viable cells seeded (Fig.2A). The adherent MSPCs from G3 and G4 were culture expanded through low density passaging and could be expanded for an average of 6.67+/-1.37 passages (G3) and 6.86+/-1.88 passages (G4) before reaching senescence (Fig.2B), although some cultures could be expanded for over 10 passages.

Whenever CFU-f’s were obtained (from G3 and G4), passage 3 cells were allowed to reach confluence and were then placed in human Mesencult adipogenic, osteogenic, or chondrogenic medium (StemCell Technologies), n = 5 for G3 and n = 6 for G4. All primary MSPC cell line tested were shown to be able to differentiate into adipocytes, osteoblasts, and chondrocytes in vitro (Fig.2C). Taken together these results indicate that the hip pulvinar is not an important stem/progenitor cell reservoir in healthy patients but is an important source of MSPCs in the osteoarthritic pulvinar.

### Effects of age and OA on the pulvinar’s immune landscape

We isolated total RNA from three patients from each of the four groups (12 patients in total) and performed a probe-based hybridization assay using the Nanostring Immunology Panel V2, capable of detecting up to 594 immune-related human genes. Through analysis with the Rosalind platform, we observed a high correlation between samples from patients without OA that patients with OA highlighting that the technology was able to report similarities between the samples of the same group as well as differences with other groups (Fig.S1A). Cell type analysis using the Advanced nSolver analysis algorithm integrated within the Rosalind platform revealed a notable decrease in the overall number of CD45+ cells with the onset of OA (Fig.S1C), suggesting that there is a significant decrease immune infiltration of the ssAT with the onset of OA. Similarly, neutrophils and macrophages were also notably attenuated with the onset of osteoarthritis (OA), alongside an increase in exhausted CD8+ cells (Fig.S1C). Conversely, dendritic cells (DC) and cytotoxic cells did not follow a specific pattern, showing perhaps patient-patient variability. These immunological perturbations, encapsulated in our cell type analysis, suggest a complex interplay between the innate and adaptive arms of the immune system at the advent of OA. The reduction in innate immune cells could imply an impaired clearance capacity, allowing for the accumulation of stimuli that drive T-cell exhaustion. On the other hand, the enrichment of exhausted CD8+ T cells may reflect an adaptive immune response that has been chronically engaged and is now in a state of functional decline.

### OA-specific changes in gene expression

Using the Nanostring mRNA data, we next wanted to deconvolute the effects of normal aging from those specifically attributed to OA onset, therefore, we ran the following pairwise comparisons: (G3 & G4 vs G1 & G2) and (G3 vs G2) between OA and non-OA groups to identify differentially expressed genes (DEGs, Fig3A). We subsequently established cutoff values of 1.25 and −1.25 for upregulated and downregulated genes, respectively, along with a p-value threshold of 0.05, to identify genes that were significantly upregulated or downregulated across each comparison. The results of the DEG analysis are summarized in (Fig.5). Notably, seven genes were observed to be upregulated concurrently in both comparisons, including CDH5 (VE-cadherin), THY1 (CD90) and NOTCH1. Intriguingly, these genes have also been reported to exhibit upregulation in proliferating stem/progenitor cell populations^19–21^. In addition to these genes at the intersection of these 2 comparisons, our analysis further revealed a distinct set of genes unique to each comparison. Particularly, in the comparison between groups (G3 & G4 vs G1 & G2), unique genes such as CXCL2, IL2RB, IL6, MAPK11, MME, and PPBP were upregulated with OA. Conversely, the comparison of age-matched patients with and without OA (G3 vs G2) revealed a distinct profile with genes like CTSG, CX3CL1, ENTPD1, FCER1A, NFATC2, PECAM1 (CD31), and TNFSF10 being uniquely identified. To understand the significance of these genes, we performed a Reactome (Fig.3B, C), KEGG (Fig.S2A & Table.S1) and WP (Fig.S2B & Table.S2) Enrichment Analysis using ClusterProfiler^22^ on genes exhibiting significant Log fold changes and plotted the top 8 most enriched terms as well as the genes linked to the term as heatmaps (Fig.3B & C). This analysis revealed an enrichment of several key signaling pathways differing between OA and non-OA groups. The upregulated genes exhibited a notable enrichment of IL6 signaling pathways (Fig.3B), underscoring the importance of IL-6 in OA^23^. Additionally, IFN-gamma signaling was elevated in OA, aligning with the recognized role of this cytokine in promoting cartilage degradation^24^. We additionally identified seven genes that exhibited significant downregulation in OA at the intersection of all groups (Fig.5): CCL20, IFNB1, IL1RL2, IL1RN, IRGM, KLRF1, and TNFRSF17 (Fig.3A and B, Fig.5). Among these, CCL20, IFNB1, IL1RL2 and IL1RN play important roles in modulating immune responses (Table.S1 & 2). Additional genes uniquely associated with the (G3 & G4 vs G1 & G2) comparison were also identified, including CD55, FN1, IL20, IL21, IRF5, KLRC4, KLRK1, NOD2, and PIGR (Fig.3B). These genes play significant roles in the regulation of T cell differentiation and function (Fig.S2A) and IL-20 signaling (Fig.3B & C). Moreover, in the G3 vs G2 comparison, five genes were downregulated: CD1A, CXCL11, FADD, ICOS, RAG1. The downregulation of FADD specifically is concordant with the observed downregulation in apoptosis term (Table.S4). We also noted an enrichment for IFN-alpha/beta signaling (Fig.4B), concordant with the observed enrichment of the JAK/STAT pathway, a known downstream effector of IFN-alpha/beta signaling^24^ (Table.S4). Taken together, the observed changes reveal the complexity of OA and identify several immune-related molecular pathways associated with OA.

**Figure 3.**
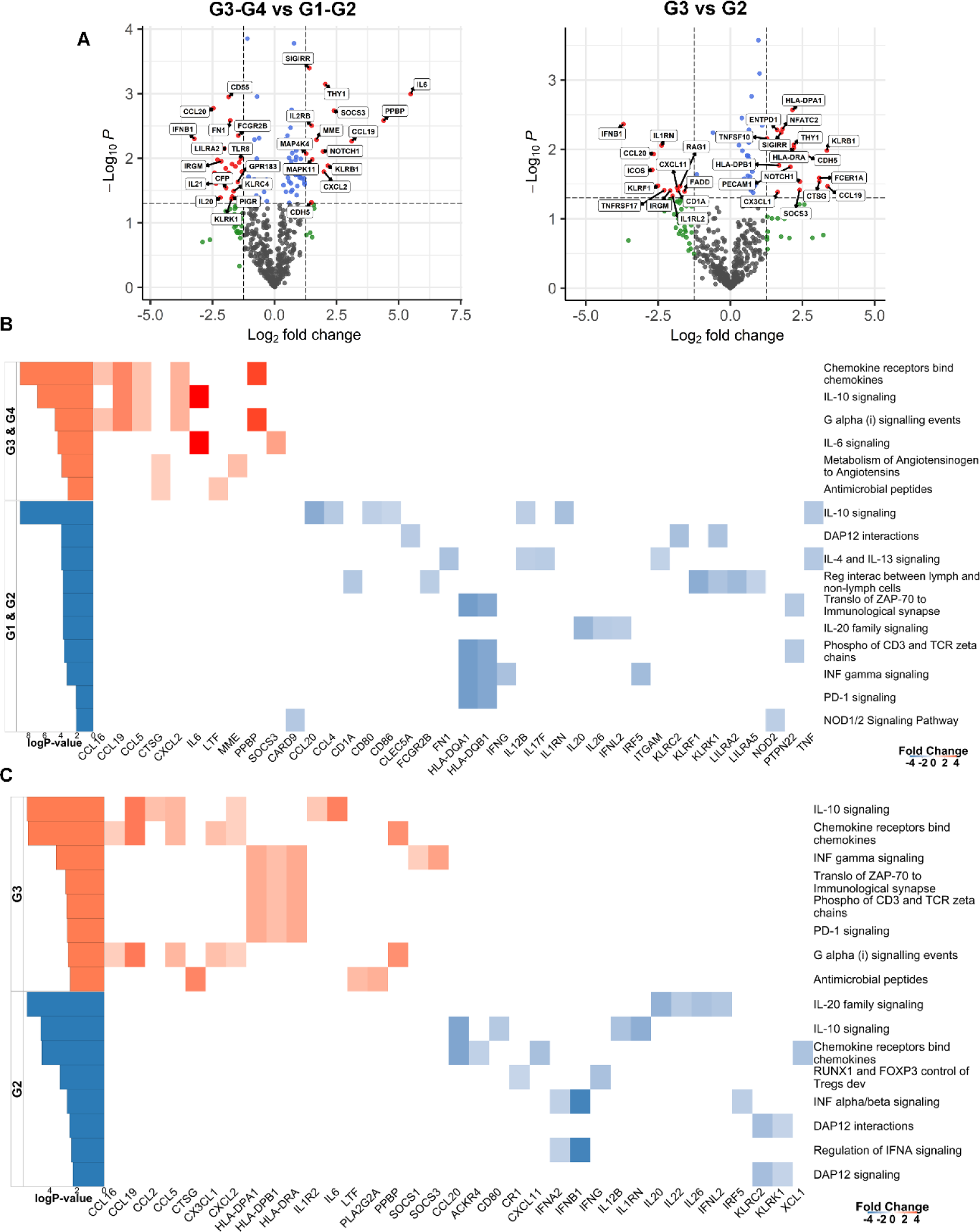
mRNA-based analysis of immune-related molecular pathways in the human osteoarthritic pulvinar. Analysis of bulk RNA samples from pulvinar samples using the Nanostring nCounter human Immunology v2 panel, analyzed using Rosalind and Reactome enrichment analysis, n=3 patients/group. Pairwise comparisons of changes in the gene expression profile between arthritic (G3 and/or G4) and the baseline non-arthritic (G1 and/or G2) patients. For all comparisons, the volcano plots shows the differential gene expression (DGE) between the condition (G3 and/or G4) and baseline (G1 and/or G2). The Reactome analysis shows the top eight immune-related molecular pathways that are differentially regulated (with a significant Log P-value) between the in the (G3 and/or G4) vs (G1 and/or G2) comparison and the differentially expressed genes identified for each pathway (with a significant LogFC). A) Volcano plots showing the differentially expressed genes in the arthritic vs non-arthritic patients. B) Comparison between the enriched terms in the arthritic groups (G3 and G4) and the baseline non-arthritic (G1 & G2). C) Age-matched comparison of the enriched terms between the arthritic patients (G3) and the baseline non-arthritic (G2). Volcano plot legend: Red: |log2FC| >1.25 and p-val<0.05, Green: |log2FC| >1.25, Blue: p-val<0.05, Grey: Insignficant

**Figure 4.**
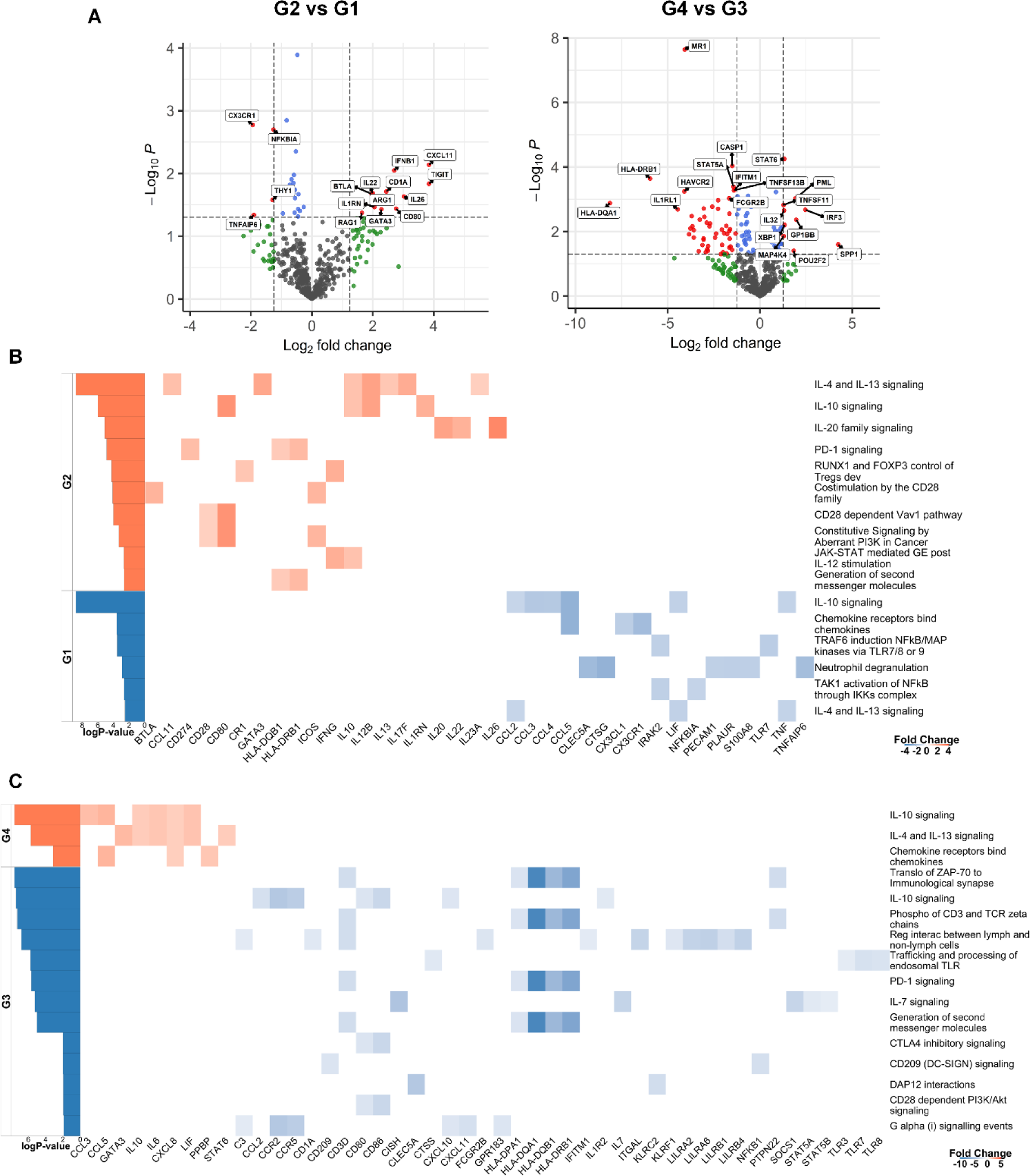
mRNA-based analysis of immune-related molecular pathways in the aging human pulvinar. Analysis of bulk RNA samples from pulvinar samples using the Nanostring nCounter human Immunology v2 panel, analyzed using Rosalind and Reactome enrichment analysis, n=3 patients/group. Pairwise comparisons of age-related changes in the gene expression profile between non-OA (G1 and G2) and OA (G3 and G4) patients. For all comparisons, the volcano plots show the differential gene expression (DGE) between the condition (G3 and/or G4) and baseline (G1 and/or G2). The Reactome analysis shows the top eight immune-related molecular pathways that are differentially regulated (with a significant Log P-value) between the condition and baseline and the differentially expressed genes identified for each pathway (with a significant LogFC). A) Volcano plots showing the differentially expressed genes with aging. B) Enriched pathways with aging on non-arthritic patients B) Enriched pathways with aging in arthritic patients. Volcano plot legend: Red: |log2FC| >1.25 and p-val<0.05, Green: |log2FC| >1.25, Blue: p-val<0.05, Grey: Insignficant.

**Figure 5.**
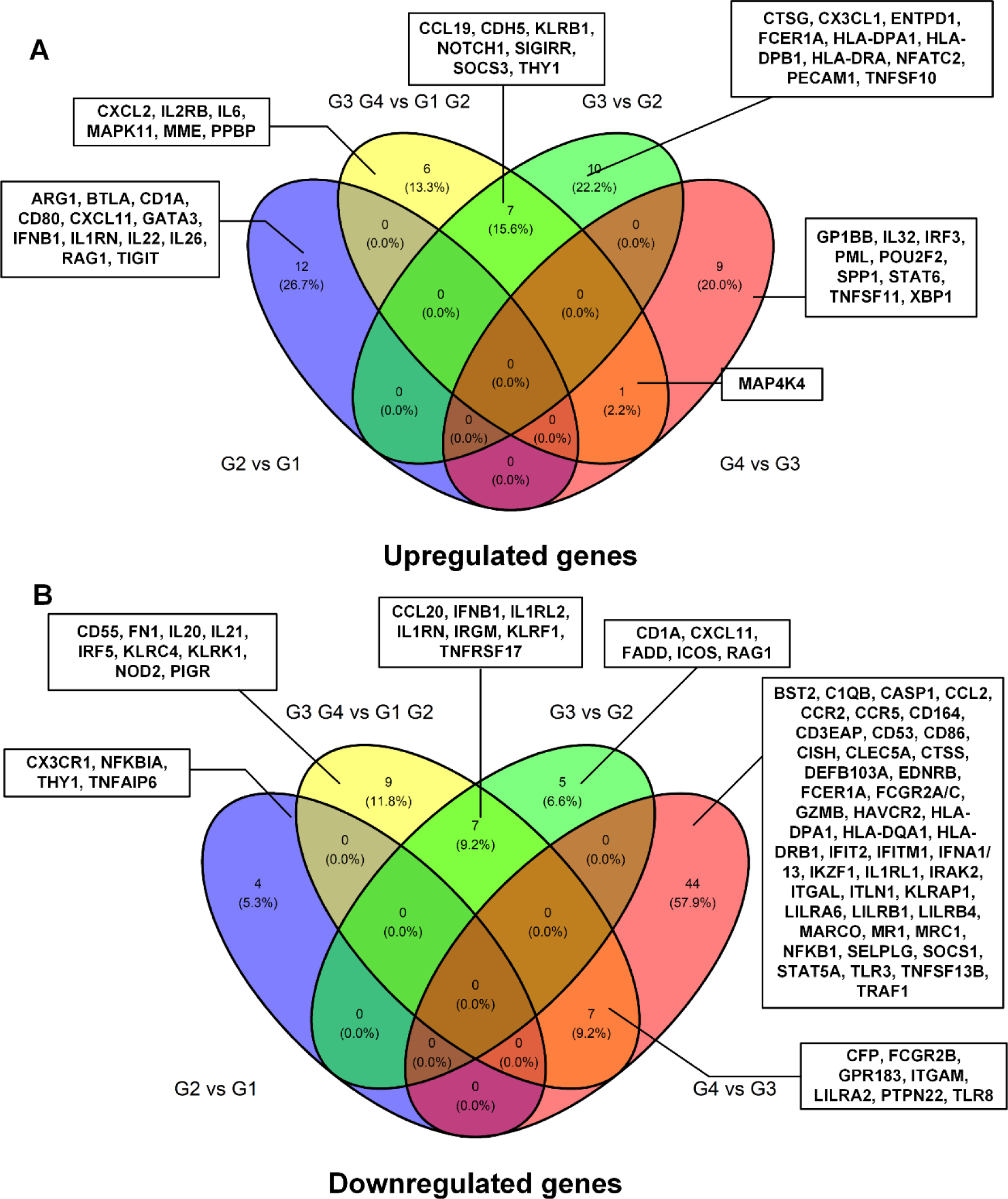
Summary of the significant DEGs in the pairwise comparisons. Venn diagram showing the number of genes shared or unique to each of the pairwise comparisons. The genes at each specific intersection have been annotated. A) Upregulated genes. B) Downregulated genes.

### Age-specific changes in gene expression

To further uncouple the effects of normal aging from those of OA, we performed pairwise comparisons based on patient age groups within the OA cohort (Fig.5). We observed that there were no common genes in the immunology panel that were upregulated in the aging process in non-OA groups (G2 vs G1) nor in the OA patients (G4 vs G3) (Fig.5). We therefore investigated DEGs within each comparison. We found an upregulation in genes such as GP1BB, IL32, IRF3, PML, POU2F2, SPP1, STAT6, TNFSF11 and XBP1 in the (G4 vs G3) comparison (Fig.5A). These genes showed an enrichment in terms related to other inflammatory conditions such as Rheumatoid arthritis and autograft rejection (Fig.S4 & S5) as well as an implication in osteoclast differentiation (Table.S5). In the non-OA cohort, we observed an upregulation of genes such as ARG1, BTLA, IFNB1, IL1RN, IL22, IL26, RAG1 and TIGIT which have shown a role in Th17 cell differentiation, autoimmune disease and proinflammatory and profibrotic disorders (Fig. S4 & Table.S6). These changes highlight the potential proinflammatory landscape of aging which may contribute to disease severity through the secretion of SASP^25,26^. By examining the downregulated genes in the G4 vs G3 comparison, we identified a list of 44 genes (Fig.5) including CASP1, NFKB1, PRF1 and IFITM1, which are enriched in apoptosis and cell cycle arrest (Table.S6 & S8). These findings may reflect age-related cellular senescence rather than direct involvement in OA especially since similar pathways were also enriched in the downregulated terms of non-OA patients (Table.S6).

### OA- and age-associated changes in the pulvinar adiposity, vascularity, and innervation

Sub-synovial ATs are thought to act as neuroendocrine organs, and to play a role in proprioception and mechano-sensing. Our gene expression analysis (Fig.3) also identified endothelial cell markers being overexpressed in OA (CDH5, PECAM1). Therefore, we next asked if the cellular architecture of the pulvinar was affected by OA and aging. To do this, we fixed the samples and sectioned them at 300μm thickness using a vibratome. We then performed immunostaining for a vascular marker (PECAM1/CD31), and peripheral neuron marker (peripherin), and counterstained the samples with a lipid dye (BODIPY) to stain adipocytes (Fig.6A). We optically cleared the thick samples and performed confocal imaging. All fluorescence channels were segmented, and distances between segmented objects were computed as well as the volume of these objects.

**Figure 6.**
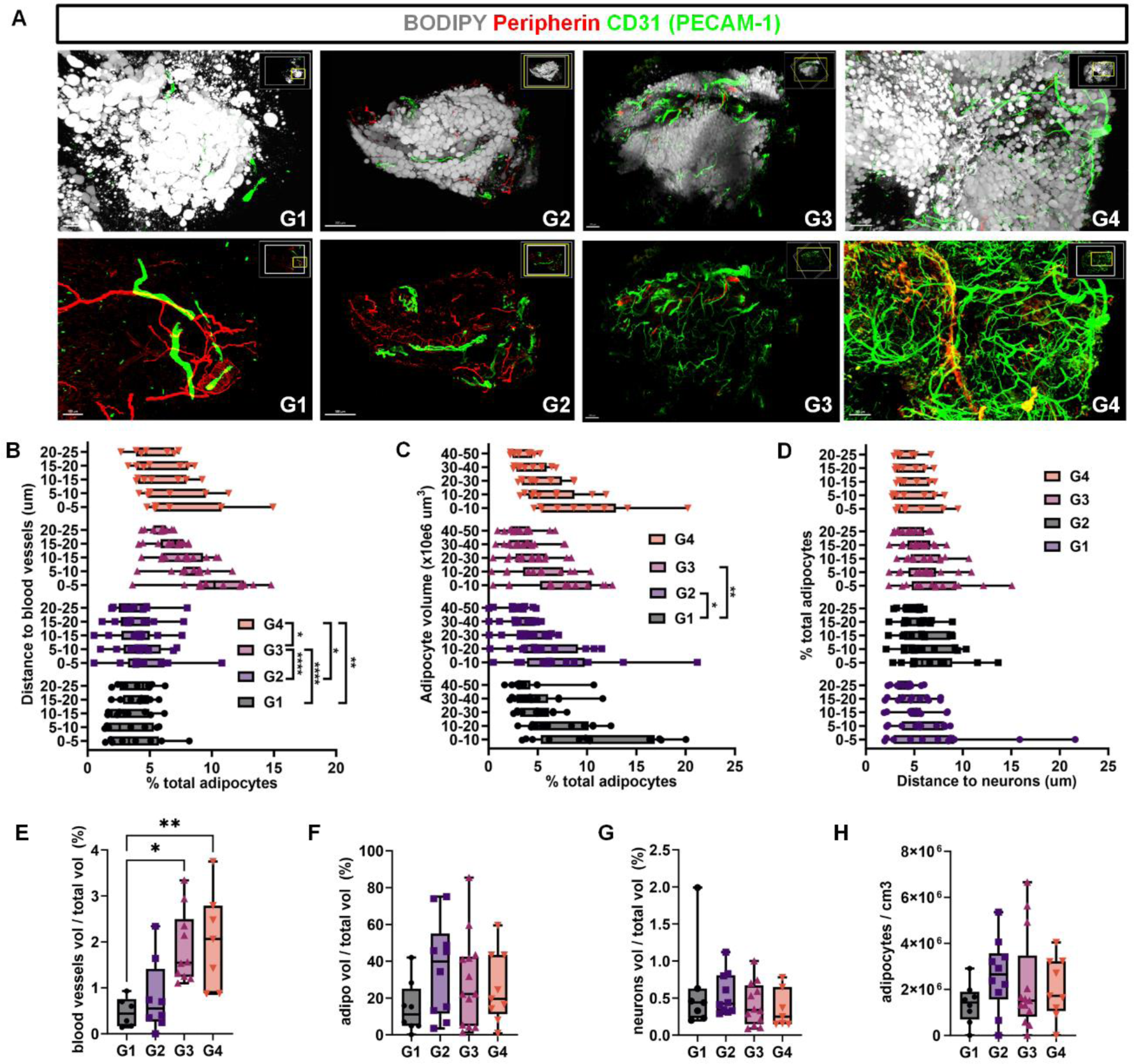
Cellular changes in the aging and osteoarthritic human hip pulvinar. Freshly obtained human hip pulvinar samples from all patient groups were fixed and stained for adipocyte lipid droplets (BODIPY), peripheral neurons (peripherin), and blood vessels (CD31, PECAM-1). Samples were then optically cleared before confocal imaging. All fluorescent channels in the images were then segmented for quantification. Two-way ANOVA was used with a significance threshold set at p<0.05. A) Representative images of samples from all groups showing all fluorescent channels (top), as well as peripherin and CD31 only (bottom). Scale bars=100μm, 300μm, 400μm, and 200μm for G1, G2, G3 and G4, respectively. B) Frequency distribution of the distance between adipocytes and blood vessels shows important age- and OA-related changes in adipocytes vascularization. C) Frequency distribution of adipocytes cellular volume shows minor changes in adipocyte volume, an indicator of adipocyte maturity. D) Frequency distribution of the distance between adipocytes and peripheral neurons shows only minor, mostly age-related changes in adipocytes innervation. E-H) Total tissue vascularity (E), adiposity (F), and innervation (G) was quantified as well as the total number of adipocytes per tissue volume unit (cm^3^) (H). We observed a significant OA-related increase in tissue vascularity (E), and small but not statistically significant age- or OA-related variations in tissue adiposity (F) and number of adipocytes per tissue volume unit (H).

When we compared the distance between adipocytes and blood vessels across the groups (Fig.6B), we observed significant differences between nearly all groups (except between G1 and G2). Most notably, in OA samples we observed significantly more adipocytes in direct contact with blood vessels (0-5μm bins), with up to 10.4 +/- 1.5% adipocytes directly touching a blood vessel in G3. We also observed that the adipocytes from younger patients (G1) were significantly smaller (presumably less mature) than those from G2 and G3 (Fig.6C). However, the distance between adipocytes and peripheral neurons was not significantly different between groups (Fig.6D). Intriguingly, up to 7.4 +/- 1.7% of the adipocytes were directly innervated by neurons (0-5μm bin in G3), suggesting that ssATs may indeed act as neuroendocrine organs.

Similarly, when we calculated the total volume of blood vessels within the tissues, we observed an increased vascularity in the ssATs from OA patients. In non-OA patients, the bloods vessels accounted for 0.47 +/- 0.16% and 0.95 +/- 0.40% of total tissue volume for G1 and G2 respectively, but for up to 2.04 +/-0.39% and 2.14 +/-0.53% of total tissue volume in G3 and G4, respectively (Fig.6E). The volume of total adipocytes (Fig.6F), neurons (Fig.6G) and the number of adipocytes per cm^3^ of tissue did not differ between the groups. In summary, these results indicate that OA is associated with a significant increase in blood vessels density in the hip pulvinar and that a significant number of ssAT adipocytes are directly associated with blood vessels and neurons.

## Discussion

OA is a complex disease implicating signaling from various tissues of the joint. The ssAT has been proposed to play a particular role as a stem cell reservoir, in proprioception and mechano-sensing, as a neuroendocrine organ, and as a modulator of inflammation^13–16^. In this study, we aimed to identify the various cellular and molecular changes in the ssAT with the onset of OA and aging and particularly to deconvolute the effects of OA onset and progression from the effects of aging. Our findings illustrate these intricate roles of ssATs in joint homeostasis and repair.

Central to our findings was the increase in tri-potent mesenchymal stem/progenitor cells (MSPCs) concomitant with OA onset. This observation contributes to the understanding of the dynamics of mesenchymal stem/progenitor cells (MSPCs) in osteoarthritis (OA). Notably, the elevated presence of MSPCs may be attributed to several factors: the potential migration of these cells from surrounding tissues into the joint^27^, the possibility of resident cells within the sub-synovial adipose tissue (ssAT) to dedifferentiate to an MSPC-like state in response to OA^28^, or a significant activation of the MSPCs with OA^29,30^ that is otherwise undetectable in non-OA conditions. This last point echoes findings from previous studies which have identified increased numbers of MSPCs in the synovial fluid of OA patients, suggesting that the increased abundance of these cells is a hallmark of the joint’s response to OA^31,32^. This pronounced expansion in OA could be linked to the increased tissue degeneration which requires a surge in reparative cells, that are however unable to overcome the degenerative process. Our transcriptomic analysis via Nanostring technology revealed that tissues from OA patients exhibit increased expression of markers like DPP4 and THY1, aligning with literature reports on their upregulation during joint repair^21,29,30^. These markers are less expressed in healthy ssAT, suggesting that the activation and subsequent proliferation of MSPCs are specific responses to OA pathogenesis, reflecting an innate repair mechanism that becomes activated during the disease.

The immune landscape, altered by aging and disease, exhibits a reduction in innate immune cells, such as neutrophils and macrophages, alongside an increase in exhausted CD8+ cells in OA patients. This is also particularly noticed in the enrichment of terms linked to the adaptive immune system in OA (T-cell activation terms) and the enrichment of innate immune system terms in non-OA patients (Toll-like receptor signaling). This shift paints a picture of an immune system wrestling with chronic inflammation and potentially hampered in its response to continuous joint degradation. Notably, genes linked to the presence of pro-inflammatory macrophages and fibroblast activation, such as IL6, IFN-gamma, and THY1, are upregulated in OA^29,33,34^, emphasizing their significance in promoting the systems responsible for clearing up damage. Additionally, the role of adaptive immunity, particularly the observed increase in exhausted CD8+ T cells and enrichment in pathways linked to T helper cell activation and differentiation (Th1, Th2, Th17), along with regulatory T cells (Tregs), is crucial in OA^35,36^, orchestrating a delicate balance between pro-inflammatory and anti-inflammatory signals that influences the disease’s outcome. Th1 and Th17 cells tend to promote inflammation and joint degradation, while Th2 and Tregs serve to temper the inflammatory response^36^.

In comparing OA with non-OA patients, we noted an upregulation in genes associated with inflammation (IL6, MME)^34,37,38^, immune regulation (SIGIRR, THY1, SOCS3, CCL19)^35,39^, and tissue remodeling (NOTCH1, THY1, MME)^19,29,37^. Conversely, genes involved in complement activation, cytokine signaling, and immune recognition were downregulated. The prominence of IL6 and THY1 upregulation supports their proposed roles in OA pathology. Specifically, IL-6 acts through pro-inflammatory that contributes to cartilage degeneration in OA^34^. THY1 on the other hand marks fibroblast activation pathways^29^. The later has been shown to be important in the development of scaring tissues in the synovium, ligaments and articular cartilage in OA^40,41^ but our findings suggest ssAT might play a role in promoting scaring and joint stiffness as well. ^31^ Additionally, the elevated IFN-gamma signaling in OA can be contrasted with an enrichment for IFN-beta signaling in non-OA conditions, suggesting the significantly different roles these two players can play in promoting the progression of the disease^24^ and promoting joint health^42^, respectively.

Furthermore, we observed upregulation of NFATC2 and ENTPD1—important for T-cell activation and inflammation modulation, respectively^43,44^. The upregulation of NFATC2 not only influences T-cell activation but also intersects with cartilage integrity and protection mechanisms^45^. Indeed, since the samples were not enriched for CD45+ immune cells, the upregulation of these genes could be a hallmark of their upregulation in other cell types. Particularly, the suppression of NFATC1 and NFATC2 in chondrocytes of the superficial layer of the articular cartilage has been shown to have a protective role against spontaneous OA^45^. Conversely, ENTPD1 suppression in chondrocytes has been suggested to offer protection against OA by decreasing the release of NO, MMP-13 and MMP-3^46^. Therefore, the increased expression of NFATC2 and ENTPD1 observed in our findings could suggest their respective roles in promoting cartilage damage for NFATC2 and a rescue attempt by ENTPD1.

Our analysis extended to discerning the effects of aging on the ssAT, revealing distinct gene expression patterns that suggest aging and OA alter gene expression in a different manner. Upregulation of genes like SPP1 and STAT6, involved in osteogenesis and bone remodeling, pointed to a potential aging role rather than an OA-related expression pattern^47,48^. Aging in both OA and healthy groups revealed an upregulation of pro-inflammatory and profibrotic genes, while significantly downregulated genes such as CXCR1, TNFAIP6, CASP1, and IFITM1, were linked to apoptosis and cell cycle arrest^49^. This indicates a shift in cellular senescence which could be linked to the fibrosis events observed in OA, potentially promoting the disease^49^.

Reactome Enrichment Analysis further delineated downregulated genes in pathways critical for T-cell activation, cytokine response, and immune regulation in older individuals, suggesting an aging-related shift towards enhanced immune activation, potentially aggravating OA progression. Interestingly, previous literature regarding inflammaging reports on an enhanced immune response driven by IL-6 that promotes a macrophage driven response rather than an adaptive immune system response^50^. Conversely, the enrichment in downregulated pathways associated with immune regulation in younger patients points towards a diminished inflammatory response, offering insights into the nuanced interplay between aging, immune activation, and OA progression.

Moreover, the onset of OA and aging are associated with increased tissue and adipocyte vascularization as shown by our IHC data and further highlighted by elevated expression of endothelial cell markers PECAM-1 (CD31) and CDH5 (VE-cadherin). Indeed, the significant increase in tissue vascularization with OA has been reported in the past as a hallmark of tissue inflammation, particularly, synovitis^51^. Our findings showcase furthermore how the alteration of subsynovial component to the joint could have a profound effect on the inflammatory milieu. Indeed, the direct increase in adipocyte vascularization could be an effect of the whole tissue vascularization but could also be linked to the role of adipocytes in inflammation especially in the context of obesity and metabolic syndrome consistent with previous reports of adipocytes increasing tissue vascularization^52^. These findings agree with other studies suggesting the role of angiogenesis in enhancing immune infiltration and exacerbating local inflammation^53^, thereby contributing to OA-associated pain sensitivity. Our imaging study also indicated increased vascularity of ssATs in OA patients.

Contrastingly, the consistent distance between adipocytes and peripheral neurons across all groups, along with the notable innervation of adipocytes, supports the hypothesis that ssATs might engage in neuroendocrine signaling^16^, a function that does not appear to be disrupted by OA or aging. This indicates that despite the pathological changes in the tissue architecture, the fundamental neuroendocrine function of the ssATs might remain preserved and could be altered by an increase in markers of neuroinflammation such as CGRP and SubP^54,55^. Our results contrast with a study that found decreased innervation in the osteoarthritic knee joint^56^, and it remains possible that the hip joint differs from the knee joint with regards to the impact of OA on joint innervation. The innervation of the knee joint has been well known and described for over 150 years and consists mainly of CGRP+ sensory neurons and tyrosine hydroxylase sympathetic neurons^56,57^. However, much less is known about the innervation of the hip joint^58^.

It is imperative to consider that while the density of blood vessels and proximity to adipocytes appears to be altered in OA, the overall quantity of adipocytes, neurons, and their volumetric presence did not differ significantly across the groups. This might suggest that it is not the quantity but rather the spatial organization and the secreted factors between these components that is critical in the pathophysiology of OA.

Taken together, our study captures the complexity of OA and uncouples the effects of OA from those of normal aging processes, revealing significant associations between the disease’s pathology and the vascular, immune, and cellular architecture of the hip pulvinar. By highlighting the distinct roles of inflammation, immune regulation, and stem cell activity, we identify new avenues for therapeutic intervention aimed at modulating these key aspects of OA. These findings deepen our comprehension of OA as a disease modulated by both aging and the intrinsic responses of joint tissues, opening the door to targeted strategies that address these converging pathways.

## Limitations of the study

This research represents a significant step forward in elucidating the complex interplay of cellular and molecular mechanisms underpinning osteoarthritis (OA), particularly highlighting the roles of mesenchymal stem/progenitor cells (MSPCs), vascular changes, and the immune landscape in the disease’s progression. By providing a deeper understanding of these factors, our study lays the groundwork for future therapeutic strategies aimed at targeting these key areas, potentially offering new avenues for the treatment and management of OA. Despite the significant insights garnered from our study, it’s important to acknowledge the inherent limitations that accompany our findings. Primarily, the observational nature of our study design precludes the establishment of causality between the observed cellular and molecular changes and the onset or progression of osteoarthritis (OA). Furthermore, our reliance on specific markers, such as THY1 and DPP4, to identify and characterize mesenchymal stem/progenitor cells (MSPCs) may not capture the full diversity and functional capacity of these cells within the OA and aging context. The complexity of the immune landscape in OA also presents a challenge, as the study’s scope may not encompass all the nuanced interactions between immune cells, signaling molecules, and the affected joint tissues. Finally, while our analysis provides valuable insights into gene expression changes associated with OA, it’s crucial to recognize that gene expression patterns alone may not fully elucidate the functional implications of these changes without further in-depth functional studies. Addressing these limitations in future research will be vital for advancing our understanding of OA and developing more effective therapeutic interventions.

## Methods

### Sample collection and processing

This is an IRB-approved study (CRRF ID: 2383). IAAT samples from the hips’ acetabular fossa were obtained from 135 consenting patients that underwent hip surgery (inclusion criteria included BMI <35 and the absence of inflammatory arthropathy or avascular necrosis). Patients were stratified into one of four groups: (1) young patients (<20 years), without OA undergoing hip arthroscopy; (2) young patients (20 – 40 years), without OA undergoing hip arthroscopy; (3) young patients (<40 years) with OA undergoing arthroplasty; and (4) old patients (>70 years) with OA undergoing arthroplasty. Samples were processed as follow: digested for colony assay and differentiation, immunostaining or RNA isolation.

### Cell isolation

Tissue dissociation was performed as described^59^. A specially formulated enzymatic mixture was utilized, consisting of Phosphate-Buffered Saline (PBS) supplemented with 2% Fetal Bovine Serum (FBS), 2.5 mg/mL collagenase I, 0.7 mg/mL collagenase II, 1 U/mL dispase (all enzymes sourced from Worthington), and 5µM calcium ions (Ca2+) (Sigma-Aldrich). Following preparation, the samples underwent agitation on a shaker at 37°C for 45 minutes to facilitate cell separation. Subsequent to this incubation, the cell suspension was passed through a 100µm cell strainer to remove aggregates, and the filtered cells were seeded into cell culture dishes at different densities for colony count and at 50,000 cell/cm2 for cell expansion and differentiation in Human MesenCult proliferation Medium (StemCell Technologies).

### In Vitro differentiation

MPSCs were passaged biweekly or when they reached 80% confluency, until passage 2. Then, they were harvested and plated in six wells of a 24 well plate for differentiation assays at a cell density of 10000 cells/cm2 in MesenCult media. Upon reaching 80% confluency, the MesenCult media was replaced with adipogenic differentiation media or osteogenic differentiation media (StemCell Technologies), for 10 days. This was followed by a brief fixation for 15min and staining with Bodipy (2uM) and DAPI (1:500 dilution of a 2mg/mL stock solution) for the adipogenic wells and Alizarin red for the osteogenic wells. For the chondrogenic differentiation, cells were pelleted in Chondrogenic differentiation media (StemCell Technologies) and media was replaced as recommended for 21 days followed by fixation and staining with Sox9, DAPI and Collagen I.

### Tissue sectioning

Adipose tissue was processed as described^60^: tissue from patients undergoing surgery was fixed in 4% paraformaldehyde at 4℃ for 16h. Tissue was then embedded in 3% agarose (high gel strength grade, Bioshop, CAS#9012-36-6) and sectioned to 300um thick sections.

### Immunostaining

Immunostaining of ssAT sections was prepared as published^60^. Sections were placed on Superfrost microscope slides using silicone spacers (Grace Biolabs). Sections were then blocked and permeabilized using a blocking buffer comprised of 0.1M Tris, 0.15M NaCl (pH 7.5), 0.05% Tween-20, 20% dimethyl sulfoxide (DMSO), 5% donkey serum, and 0.3% Triton X-100, all procured from Sigma-Aldrich, for 1 hour at room temperature. Following blocking, sections were incubated with primary antibodies as detailed in Table.S9 in blocking buffer overnight. After primary antibody incubation, sections were washed five times for 60 minutes each in Tris-buffered saline (TBS) containing 0.05% Tween-20 and 20% DMSO. This was followed by staining with AlexaFluor-conjugated secondary antibodies (at a dilution of 1:200) and counterstaining with 4’,6-diamidino-2-phenylindole (DAPI) (1:500 dilution of a 2mg/mL stock solution) and Bodipy (1:300 dilution of a 3.8mM stock solution) in blocking buffer, also overnight at RT. The fluorophores utilized were AlexaFluor 488 and 633. Following secondary antibody incubation, sections underwent five additional 60-minute washes as before. All steps were performed with gentle shaking at room temperature.

### Optical Clearing and Mounting of Sections

To enhance optical clarity and permit deeper imaging, sections were optically cleared using a modified Ce3D protocol^61^. The clearing medium was prepared by dissolving histodenz to 88% in TBS, supplemented with 0.1% Tween-20 and 0.01% sodium azide (NaN3), and adjusting the pH to 8.5. The refractive index of the mounting medium was adjusted to 1.467 using histodenz, as measured by a handheld refractometer (Atago). Sections were incubated in clearing medium overnight at room temperature with gentle agitation. Clearing medium was then changed and sections were mounted in the same medium with size 1.5 coverslips.

### Image acquisition and analysis

Image acquisition was conducted on a Leica TCS SP8 confocal microscope, leveraging its multiple photomultiplier tubes (PMTs). Images were captured at a resolution of 1,024 × 1,024 pixels in 8-bit format, utilizing a 1.0× zoom, bidirectional scanning mode, and a z-spacing of 2.5 µm. For the analysis, images underwent lossless compression and were visualized and segmented using Imaris software (version 9.9, Bitplane). Bodipy, CD31 and Peripherin were then segmented using the surface segmentation function and statistics were then exported, binned using R and reexported for plotting using GraphPad Prism version 9. Data was then plotted as frequency distribution, and statistical significance was assessed using a multiple comparison test and two-way ANOVA, with a threshold set at p < 0.05. Specific n and p values reported in the text or figure legends are indicated where applicable.

### RNA isolation

RNA isolation was executed utilizing a refined Trizol method paired with on-column purification for enhanced RNA recovery. Initially, samples were treated with 500 μL of phenol solution per 50 mg of tissue, followed by mechanical disruption via homogenization. An initial 5min centrifugation at 4℃ allowed for the separation of the mixture into 3 layers: fat, RNA isolate, and pellet phases. The RNA phase was carefully extracted, minimizing lipid contamination. Subsequent chloroform addition (400 μL per sample) and phase separation further refined the RNA isolation. Using the Norgen#61000 clean-up kit, RNA was precipitated with an equal volume of 70% ethanol, filtered through a spin column, and washed to remove impurities. The purified RNA was then eluted in 8-15 μL, depending on desired concentration. Storage conditions were −20°C for short-term and −70°C for long-term.

### Nanostring assay

To evaluate gene expression profiles within subcutaneous adipose tissue (ssAT) samples, we employed the NanoString nCounter technology, focusing on immune signaling pathways. Initially, RNA integrity and quality were rigorously assessed using the Advanced Analytical Technologies, Inc. (AATI) Fragment Analyzer, ensuring that only high-quality RNA samples proceeded to analysis. RNA concentration was accurately quantified utilizing the Qubit 3.0 Fluorometer (Thermo Fisher Scientific) at the stem core facility (OHRI), adhering to stringent quality control criteria deemed optimal for NanoString assay compatibility. Following quality assessment, RNA samples were diluted to a final concentration of 300ng/5 μL. For the hybridization process, the NanoString Immunology Panel V2 (NanoString Technologies, Cat. No. XT-CSO-HIM2-12) was used. This codeset, designed to capture a comprehensive array of immune signaling genes, was mixed with the prepared RNA samples. The RNA-codeset mixture was then incubated at 65°C for 18 hours, facilitating the formation of stable RNA-reporter complexes necessary for accurate gene expression analysis. Post-hybridization, samples were processed using the nCounter MAX Analysis System (NanoString Technologies), operated under high-sensitivity settings to ensure the detection of low-abundance transcripts. The quality and integrity of the generated data were rigorously evaluated using the nSolver analysis software (NanoString Technologies), with all samples meeting or surpassing the predefined quality control benchmarks. This meticulous approach ensured that the resultant dataset was of the highest quality, enabling reliable and insightful analysis of immune signaling pathways within the ssAT samples.

### Data processing: ROSALIND

The analysis of data was conducted using ROSALIND®’s HyperScale architecture, a proprietary technology developed by ROSALIND, Inc., based in San Diego, CA. The Quality Control (QC) phase included the generation of Read Distribution percentages, violin plots, identity heatmaps, and Multidimensional Scaling (MDS) plots. The process of normalization, as well as the determination of fold changes and p-values, adhered to the guidelines set forth by Nanostring. Specifically, ROSALIND® utilizes the nCounter® Advanced Analysis protocol, which involves normalizing counts within a lane by the geometric mean of the lane’s normalizer probes. The selection of housekeeping probes for normalization is carried out through the geNorm algorithm, as implemented within the NormqPCR R library^62^. Additionally, ROSALIND calculates the abundance of various cell populations utilizing Nanostring’s Cell Type Profiling Module, and filters the Cell Type Profiling outcomes to include only those results with a p-Value of 0.05 or greater. The calculation of fold changes and p-values is executed via the fast method detailed in the nCounter® Advanced Analysis 2.0 User Manual, with p-value adjustments being made through the Benjamini-Hochberg method for estimating false discovery rates (FDR).

### Go term, Wikipathway and KEGG enrichment analysis

Following the identification of differentially expressed genes (DEGs) using the Rosalind platform, we conducted an enrichment analysis to uncover the biological pathways and processes associated with these genes. This analysis was executed using the clusterProfiler^22^ package in R. Initially, we focused on genes exhibiting a log fold change greater or equal to 1.25 and less or equal to −1.25, intentionally disregarding the p-value at this stage to enrich for biological terms and pathways, thereby enhancing the comprehensiveness of our analysis. Subsequently, to refine our results and focus on statistically significant associations, we excluded terms with a p-value greater than 0.5. This filtering step ensured that only biologically and statistically significant pathways and processes were considered in our subsequent analyses. We specifically obtained information related to Reactome pathways^63^, KEGG pathways^64^ and WikiPathways^65^ terms. This comprehensive approach allowed us to capture a wide spectrum of biological insights associated with the DEGs identified in our study. For the visualization of our enrichment analysis results, we employed two R packages: **ComplexHeatmap** and **EnhancedVolcano**.

## Notes

### Competing Interest Statement

The authors have declared no competing interest.

