## Supplementary material for "Cellular and molecular changes in the human osteoarthritic and aging hip pulvinar": Fig.S

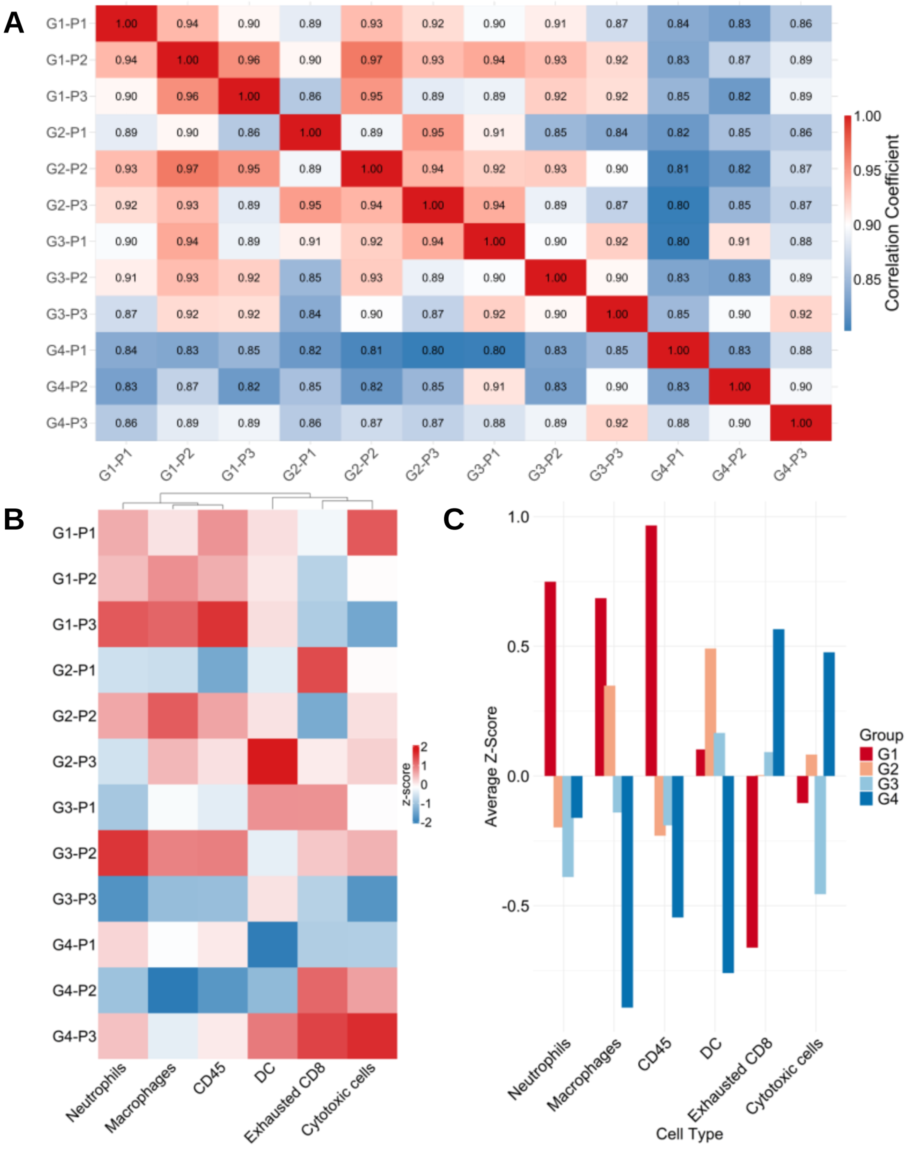


**Figure S1. mRNA-based analysis of immune cell infiltrates in the human hip pulvinar.** Analysis of bulk RNA samples from pulvinar samples using the Nanostring nCounter human Immunology v2 panel, analyzed using Rosalind, n=3 patients (P)/group. A) Correlation matrix showing patient to patient correlations. A higher variability was observed in older osteoarthritic patients (G4, P1-3). B) Heatmap of individual patient z-scores for the various immune cell types identified. Patients were not clustered based on their individual z-scores across the matrix but by experimental group. C) Barplot of average z-score per group, showing the relative changes in abundance of immune cell types in an age and OA-dependent manner.

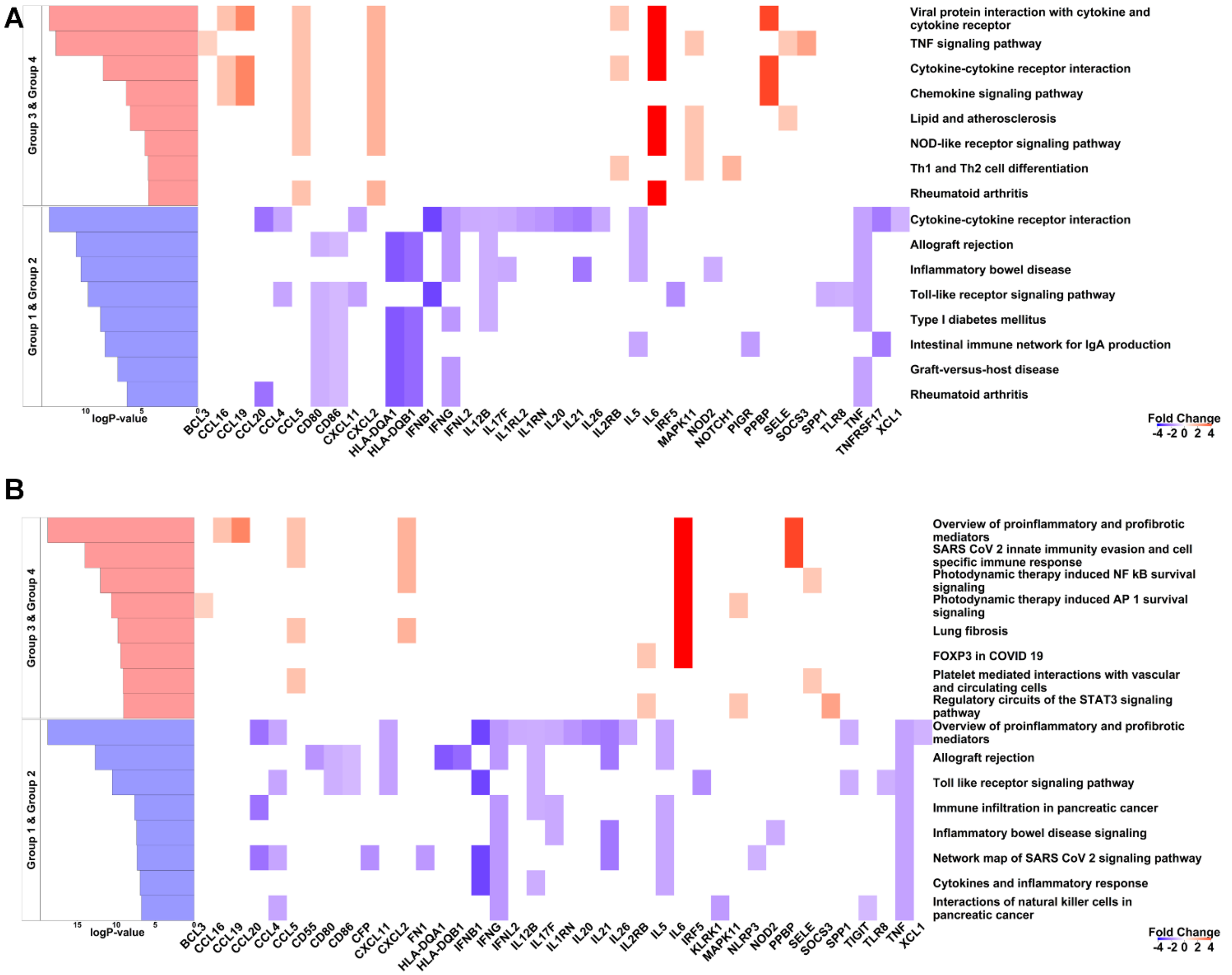


**Figure S2. Enriched pathways with the onset of OA in the (G3 & G4) vs (G1 & G2) comparison.** A) Top eight enriched KEGG pathways. B) Top eight enriched Wikipathways.

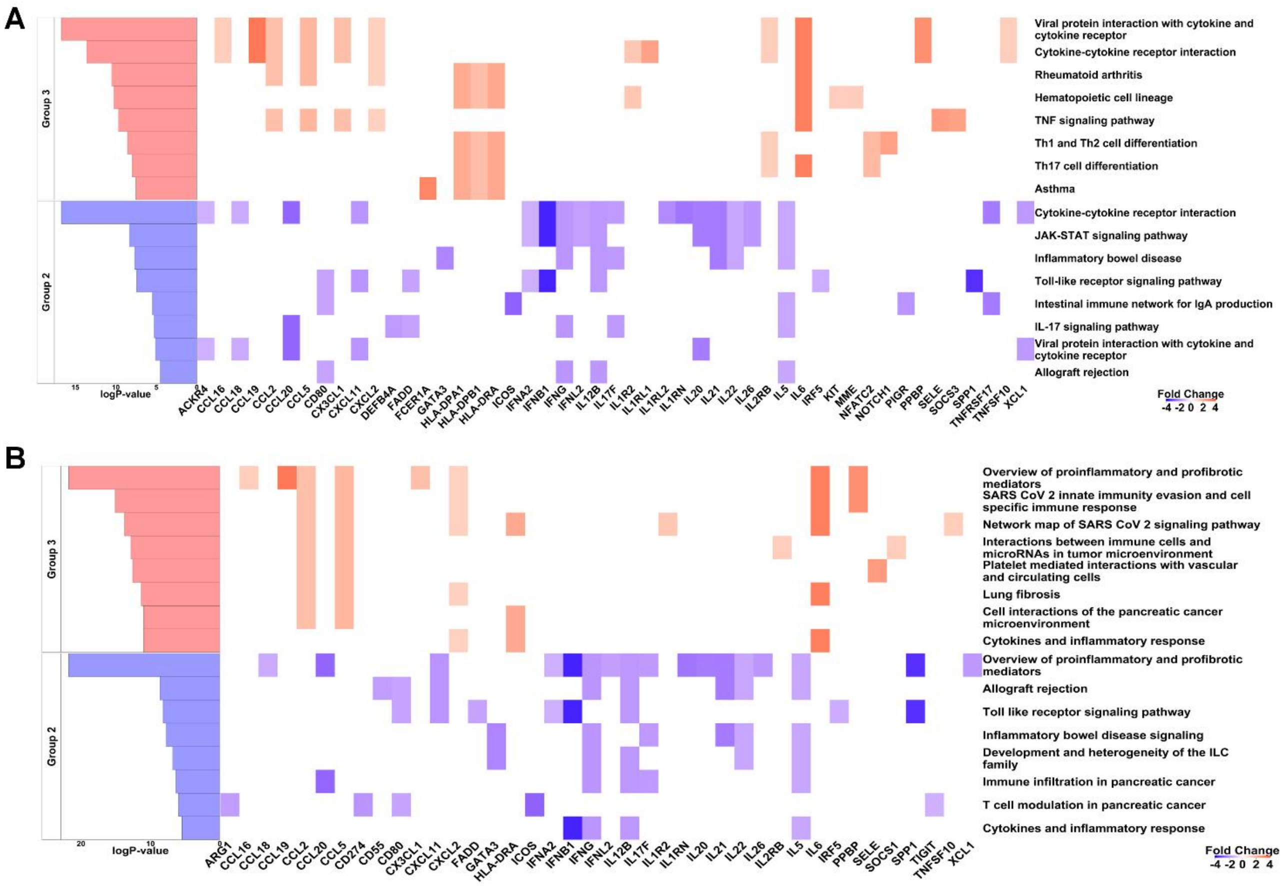


**Figure S3. Enriched pathways with the onset of OA in the G3 vs G2 comparison.** A) Top eight enriched KEGG pathways. B) Top eight enriched Wikipathways.

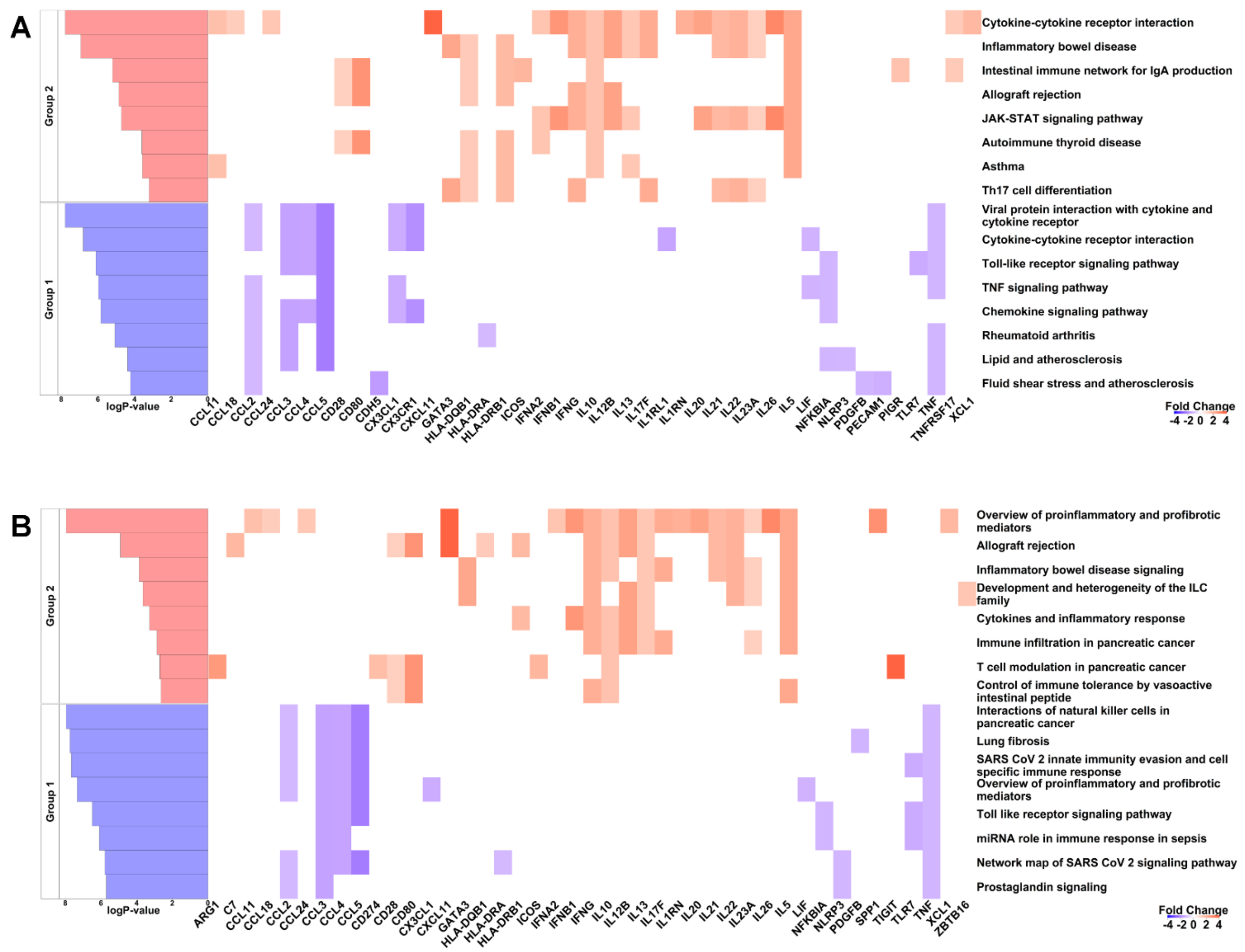


**Figure S4. Enriched pathways with the onset of OA in the G2 vs G1 comparison.** A) Top eight enriched KEGG pathways. B) Top eight enriched Wikipathways.

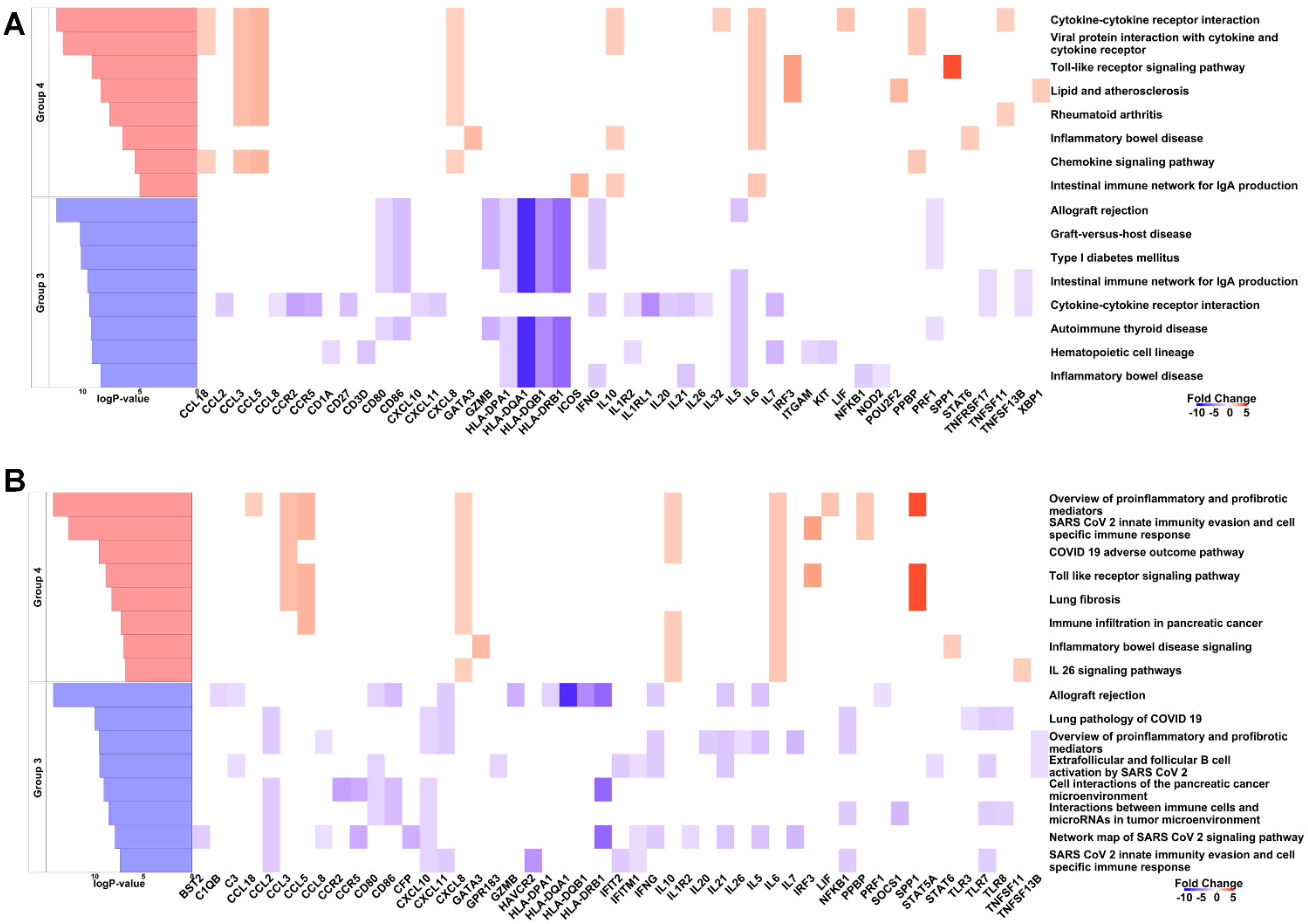


**Figure S5. Enriched pathways with the onset of OA in the G4 vs G3 comparison.** A) Top eight enriched KEGG pathways. B) Top eight enriched Wikipathways.

### Table S9. Antibodies used

| Catalog number | Target | Alternative names | Concentration | Company | Target Species | Host Species |
|---|---|---|---|---|---|---|
| 17399-1-AP | Peripherin | PRPH, PRPH1 | 1:50 | Thermofisher | Human, Mouse, Rat | rabbit |
| D3922 | Bodipy (3.8mM) | | 1:300 | Invitrogen | | |
| AF806 | CD31 | PECAM1 | 1:50 | R&D Biotechne | Human | goat |
| pa185319 | Col1a1 | Collagen 1a1 | 1:50 | Invitrogen | Mouse | rabbit |
| AF3075 | hSOX9 | | 1:50 | R&D systems | human, mouse | goat |
